# Evaluation of the psychometric properties of the HIV Disability Questionnaire among adults living with HIV in the United Kingdom

**DOI:** 10.1101/556183

**Authors:** Darren Brown, Bryony Simmons, Marta Boffito, Rachel Aubry, Nneka Nwokolo, Richard Harding, Kelly O’Brien

## Abstract

**Objective:** To evaluate the psychometric properties of the HIV Disability Questionnaire (HDQ) among adults living with HIV in London, United Kingdom (UK).

**Methods:** This is a cross-sectional measurement study. We recruited and administered the self-reported HDQ, seven criterion measures, and a demographic questionnaire with adults living with HIV accessing HIV care. We determined median and interquartile ranges (IQR) for disability presence, severity and episodic scores (range 0-100). We calculated Cronbach’s alpha (α) Kuder-Richardson-20 (KR-20) statistics for disability and episodic scores respectively (internal consistency reliability), smallest detectable change (SDC) for each HDQ severity item and domain (precision), and tested 36 *a priori* hypotheses assessing correlations between HDQ and criterion scores (construct validity).

**Results:** Of N=243 participants, all were male, median age 40 years, 94% currently taking antiretroviral therapy, and 22% living with ≥2 concurrent health conditions. Median HDQ domain scores ranged from 0 (IQR: 0,7) (difficulties with day-to-day activities domain) to 27 (IQR: 14, 41) (uncertainty domain). Cronbach’s alpha for the HDQ severity scale ranged from 0.85 (95% Confidence Interval (CI): 0.80-0.90) in the cognitive domain to 0.93 (95%CI: 0.91-0.94) in the mental-emotional domain. The KR-20 statistic for the HDQ episodic scale ranged from 0.74 (95%CI: 0.66-0.83) in the cognitive domain to 0.91 (95%CI: 0.89-0.94) in the uncertainty domain. SDC ranged from 7.3-15.0 points on the HDQ severity scale for difficulties with day-to-day activities and cognitive symptoms domains, respectively. The majority of the construct validity hypotheses (n=30/36, 83%) were confirmed.

**Conclusions:** The HDQ possesses internal consistency reliability and construct validity with varied precision when administered to males living with HIV in London, UK. Clinicians and researchers may use the HDQ to measure the nature and extent of disability experienced by PLHIV in the UK, and to inform HIV service provision to address the health-related challenges among PLHIV.

## Background

For the 36.9 million people living with HIV (PLHIV) globally (1), effective treatment offers normal life expectancy (2). Consequently, PLHIV surviving past 50 years of age are increasing at exponential and unprecedented rates (3). In 2017, more than a third (39%) of PLHIV receiving HIV care in the UK were aged 50 years or older (4). Moreover by 2028 it is estimated over half of people in UK HIV care will be aged ≥50 years (5) with repeated patterns forecast in Europe and North America (6). As people live longer with chronic HIV infection, they are susceptible to health conditions arising from the underlying infection, potential side effects of treatments, and ageing (7), resulting in increasingly more prevalent multi-morbidity (8). Common conditions include bone and joint disorders (9,10), mental health conditions (11), cardiovascular disease (12-14), cancer (15,16), and neurocognitive decline (17,18). The presence of these conditions can create physical, mental, cognitive and social health-related challenges that are conceptualised as *disability* (19).

Disability is multi-dimensional (20) and episodic in nature (19). The *Episodic Disability Framework* in HIV defines disability as: physical, cognitive, mental and emotional symptoms and impairments, difficulties carrying out day-to-day activities, challenges to social inclusion, and uncertainty or worrying about the future (19). These can fluctuate on a daily basis and over the life course. Furthermore, these dimensions of disability can be exacerbated or alleviated by extrinsic contextual factors (e.g. social support and stigma) and intrinsic contextual factors (e.g. living strategies and personal attributes) (21).

As people live longer, disability assessment and treatment will be a critical component to HIV care. Measuring disability in the context of HIV is important for determining the prevalence and impact of disability, identifying interventions that may reduce disability, and to inform disability-inclusive programming (22). A valid and reliable patient-reported outcome measure (PROM) for PLHIV that can be used by PLHIV, community-based service organisations, and health providers, is critical to identify the nature and extent of disability experienced by PLHIV, assess the burden of disability living with HIV, and determine the effect of medical and rehabilitation interventions in mitigating disability. This knowledge could be used by clinicians, social service providers, health service commissioners, and policy makers, to help guide policy and program development and inform the allocation of health care resources to improve care, treatment and support, designed to reflect the long-term nature of HIV care (23).

Existing HIV-specific health status instruments tend to focus on impairments and do not fully capture the breadth of disability, specifically lacking items related to social inclusion and uncertainty (21). Impairment data alone is not an adequate proxy for disability, as people with the same impairment can experience different types and degrees of limitations, depending on personal and environmental factors (24). The majority of studies assessing disability among PLHIV focused on measurements of single impairments (25), providing a relatively narrow understanding of disability (26) that is insufficient in capturing the multi-dimensional nature of HIV (19,25). To our knowledge, there is no known self-reported data on disability, beyond impairments alone, experienced by PLHIV in the UK.

The HIV Disability Questionnaire (HDQ) is a new self-administered HIV-specific PROM developed to measure the presence, severity and episodic nature of disability among PLHIV (27). The HDQ is comprised of six dimensions of disability that were derived from the *Episodic Disability Framework*, a conceptual framework developed from the perspective of PLHIV to characterise the health-related challenges living with HIV (28,29). The HDQ is novel in that it is the sole HIV-specific PROM of disability (29). However, disability may vary depending on the context and region of the world in which PLHIV live (30). Therefore, it is critical to assess psychometric properties with a population and setting that is representative of the context in which questionnaires will be used (31). The HDQ demonstrated validity and reliability when used with PLHIV in Canada (32), Ireland (33), and the United States (34). Compared to these contexts, the UK has a different healthcare system (35), with more PLHIV accessing antiretroviral therapy and achieving viral suppression (4,36), rendering the relevance and applicability of the HDQ to PLHIV in other developed countries, such as the UK unknown.

Our aim was to assess the measurement properties, specifically internal consistency reliability, precision of measurement, and construct validity, of the HDQ for its ability to measure disability experienced by adults living with HIV in London, UK.

## Methods

We conducted a cross-sectional measurement study, to assess construct validity and reliability of the HDQ. We used quality criteria for assessing measurement properties of health status questionnaires to guide our methodological approach (31). We recruited adults, 18 years of age or older, living with HIV who attended an outpatient HIV clinic in central London, UK between March 2016 and May 2017. Potential participants were approached during regular clinic visits for their participation in the study. Ethical approval was obtained from the London Dulwich Research Ethics Committee and Health Research Authority (IRAS 165402) and the HIV/AIDS Research Ethics Board at the University of Toronto, Canada (Protocol #34126). A data sharing agreement was approved between St Stephen’s Clinical Research, Cicely Saunders Institute King’s College London, and the University of Toronto.

We administered the HDQ, a demographic questionnaire, and seven health status criterion measures (Patient Health Questionnaire (PHQ-9) (37), General Anxiety Disorder (GAD) Questionnaire (38), Fatigue Severity Scale (39), Wellness Thermometer (40), Epworth Sleepiness Scale (ESS) (41), Everyday Memory Questionnaire (EMQ) (42), and International HIV Dementia Scale (IHDS) (43). Participants had the option to either complete the questionnaires at their clinic visit, or take them home and return later via the post. Clinical characteristics were obtained from participants’ electronic medical records including number of years since HIV diagnosis, antiretroviral therapy use, most recent CD4 count (cells/mm^3^), viral load (cells/ml), and diagnosed concurrent health conditions.

### HIV Disability Questionnaire (HDQ)

The HDQ, English Version 10.5, 2017, is a 69 item self-administered questionnaire developed from the *Episodic Disability Framework*, through a community-academic partnership, to describe the presence, severity and episodic nature of disability experienced by PLHIV (19,27). The HDQ includes six disability domains: i) physical, ii) cognitive and, iii) mental and emotional health symptoms and impairments, iv) uncertainty, v) difficulty with day-to-day activities and vi) challenges to social inclusion, and one ‘good day/bad day’ health classification item. Participants are asked to rate the level of presence and severity of each health challenge on a given day ranging from 0 (not at all) to 4 (extreme). HDQ scores range from 0 to 100 with higher scores indicating a greater presence, severity and episodic nature of disability. The HDQ has demonstrated sensibility, validity, internal consistency reliability and test-retest reliability in samples of adults living with HIV in Canada, Ireland and the United States (32-34). Median administration time is 8-15 minutes.

We calculated disability presence, severity and episodic scores on the HDQ (44). Disability presence scores were calculated by summing the number of health challenges experienced for each domain and total HDQ and transforming them to a score out of 100. Disability severity scores were calculated by summing individual item scores from each domain and then linearly transforming them into domain disability severity scores out of 100. Episodic disability scores were calculated by summing the number of challenges identified as episodic in each domain and then transforming to a score out of 100. We summed the number of participants and proportion who completed the HDQ on a ‘good day’ or ‘bad day’ living with HIV (health classification). We computed missing response rates for the disability, episodic, and health classification sections of the HDQ accordingly. To maximise HDQ data, we performed mean (severity) or median (episodic) imputation on items with less than ≤10% missing responses. List wise deletion was performed for criterion measures with missing responses. We examined the distribution of HDQ item scores for a floor effect (defined as >15% of responses at the bottom (0) of the HDQ scale) and ceiling effect (defined as >15% of responses at the high end (4) of the HDQ scale).

### Demographic Questionnaire

Participants completed a self-reported questionnaire to capture demographic characteristics including; age (years), gender, ethnicity, nationality, sexuality, smoking status, household description, employment status, educational attainment, and whether registered with GP physician.

### Reliability - Internal Consistency

We calculated the Cronbach’s alpha (α) (severity scales) and Kuder-Richardson-20 statistics (episodic scales) for the HDQ domain scores to assess internal consistency reliability (degree to which the items within the instrument are correlated with each other) [α and KR-20 >0.8 defined as acceptable for individual patients] (47).

### Precision of Measurement

Standardised Error of Measurement (SEM) is a measure of precision of an instruments ability to estimate the true state of a concept. We used Wyrich criteria (48) to calculate the SEM for each item and domain score to determine the precision of measurement, meaning how accurate the observed HDQ score is with the participants’ *true* HDQ scores. [SEM = standard deviation*sqrt (1-Cronbach alpha)]. We then calculated the smallest detectable change (SDC) to determine the range in which we can be 95% confident that the true HDQ is within this range. [Observed score +/-1.96*SEM].

### Construct Validity

Measuring disability poses several challenges, with a wide range of disability definitions, and varying approaches to disability measurement (24,45). In the absence of a ‘gold standard’ approach to measuring disability (46), we assessed the accuracy of the HDQ by testing *a priori* hypotheses about predicted relationships between scores of measures that relate to disability (37-43) with scores of the HDQ.

We determined the extent to which the HDQ relates or does not relate to the seven criterion measures (37-43). The appropriate subscale scores of the HDQ were compared to criterion measures using correlation analysis. We tested 8 primary and 29 exploratory hypotheses theorising relationships between data collected in the HDQ and criterion measures using correlation coefficients (Pearson if scores normally distributed, Spearman if not normally distributed). Hypotheses included convergent and divergent construct validity testing based on previous construct validity assessment of the HDQ (32-34) and aimed to maximise data related to dimensions of the HDQ and subscale scores data collected from criterion measures. Correlation coefficients of | ≥ 0.30|, | ≥ 0.50| and | ≥ 0.70|, were defined as ‘weak’, ‘moderate,’ and ‘strong,’ respectively (31). We considered the HDQ to possess construct validity if results confirm at least 75% of the predetermined hypotheses (31). All data analyses were performed using SAS software version 9.4 (49).

### Sample Size

Our required sample size was estimated based on our construct validity analysis. To detect a weak correlation from our construct validity hypothesis, r=0.30, with a power of 0.80, and alpha of 0.05, we required a sample of n=85, inflated to at least 102 for an estimated 20% missing response rate at item level.

## Results

### Participant Characteristics

Of the 244 participants recruited, all but one identified as male (Table 1). We excluded the one participant who identified as female resulting in a total of 243 participants in this study. The median age of participants was 40 years (20% were ≥50 years), with a median year of diagnosis of 2012 (96% diagnosed 1996 or after). The majority were employed (87%), 94% were currently taking antiretroviral therapy, and 82% had an undetectable viral load (Table 1). Fifty-four percent (54%) of participants were living with a concurrent health condition in addition to HIV, and 22% reported living with at least two or more concurrent health conditions. The most common concurrent health condition was mental health (e.g. anxiety, depression, personality disorder, or schizophrenia).

**Table 1.** Characteristics of participants in analysis (n = 243)

| **Age (n=240)** |  |
| --- | --- |
| Median Age (years) (IQR) (Range) | 40 years (33, 48)<br>(Range: 22-67) |
| Number of participants (%) $\geq 50$ years | 48 (20.0%) |
| **Gender (n=243)** |  |
| Male | 243 (100.0%) |
| **Ethnicity (n=241)** |  |
| White | 213 (88.4%) |
| Black Caribbean or Black African | 6 (2.5%) |
| Indian, Pakistani or Chinese | 5 (2.1%) |
| Other | 17 (7.1%) |
| **Nationality (n=243)** |  |
| United Kingdom of Great Britain and Northern Ireland | 143 (58.8%) |
| European Union (e.g. France, Italy, Ireland, Spain, Poland, Greece, Portugal, Germany) | 75 (30.9%) |
| United States, North America | 8 (3.3%) |
| South America | 4 (1.6%) |
| Asia-Pacific Region | 8 (3.3%) |
| Other (Middle East, or Africa) | 5 (2.1%) |
| **Sexuality (n=243)** |  |
| Homosexual | 234 (96.3%) |
| Heterosexual, bisexual, unknown | 9 (3.7%) |
| **Smoking Status (n=237)** |  |
| Current Smoker | 58 (24.5%) |
| **Participants' household description: I am living.... (n=238)** |  |
| Alone | 83 (34.9%) |
| With a spouse or partner | 81 (34.0%) |
| With a flatmate/friend | 65 (27.3%) |
| Other (e.g. living with a child <18 years, living in an institutionalised residence or care home) | 9 (3.8%) |
| **Employment status (n=242)** |  |
| Employed: regular, occasional/part-time employment, self-employed, freelance | 210 (86.8%) |
| Not employed: Benefits (including Disability Living Allowance and Employment Support Allowance) | 12 (5.0%) |
| Not Employed: Income from savings, investments or pension | 15 (6.2%) |
| Support from spouse, parents, children, relatives, or friends | 5 (2.1%) |
| **Highest level of education completed (n=243)** |  |
| University | 177 (72.8%) |
| College/vocational training | 48 (19.8%) |
| Secondary | 18 (7.4%) |
| **Year of HIV diagnosis (n=241)** |  |
| Median Year of HIV Diagnosis (IQR)(Range) | 2012 (2007, 2014)<br>(Range: 1983-2016) |
| Diagnosed in 1996 or after | 231(95.9%) |
| **Taking antiretroviral therapy (n=242)** |  |
| Yes | 228 (94.2%) |
| **CD4 Count (cells/mm<sup>3</sup>) (n=239)** |  |
| Median (IQR) (Range) | Median: 676 (508, 875)<br>Range: 190-7545 |
| **Viral Load (n=236)** |  |
| Undetectable (<40 cells/ml) | 193 (81.8%) |
| **Registered with General Physician (GP) (n=243)** |  |
| Yes | 214 (88.1%) |
| **Concurrent Health Conditions\* (n=241)** |  |
| Diagnosed with other medical condition | 130 (53.9%) |
| Living with ≥2 concurrent health conditions | 52 (21.6%) |
| Diagnosed with concurrent mental health condition (eg: anxiety, depression, personality disorder or schizophrenia) | 20 (8.3%) |
| Diagnosed with concurrent malignancy, opportunistic infection, or immune reconstitution inflammatory syndrome | 18 (7.5%) |
| Diagnosed with concurrent hypertension | 18 (7.5%) |

IQR: Interquartile Range; \*as determined from the electronic health record.

### Data Completeness

The median number of missing responses for HDQ items was 7 (2.9%) for the presence and severity scale and 13 (5.3%) for the episodic scale. Proportion of missingness was <3% for the severity scale and <10% for the episodic scale. Rates for missing responses for each item were higher on the episodic scale, attributed to some participants skipping this item if they did not feel as if they had that specific health challenge. There were 10 missing responses (4.1%) for the ‘good day / bad day’ item on the HDQ.

### HDQ scores

HDQ item scores were not normally distributed. A floor effect was evident in all 69 HDQ items (100%) with >20% of the sample responding 0 (no challenge), and 52 of the items (75.4%) had a floor effect >40%. Floor effect was most prominent in the physical (95%), cognitive (100%), and day-to-day activities (100%) domains. A ceiling effect was not present in any of the HDQ items.

Highest disability presence score was in the uncertainty domain, followed by mental-emotional, challenges to social inclusion, physical symptoms, and cognitive symptoms. Highest disability severity score also was in the uncertainty domain, followed by challenges to social inclusion, mental-emotional and physical symptoms, and cognitive symptoms. Physical symptoms had the highest episodic score (Table 2). The number of participants who identified as completing the HDQ on a ‘good day’ living with HIV was 193 (79%).

**Table 2.** HDQ Summary Scores for Participants in the UK Sample (n=243)

| HDQ Subscale (# items) | HDQ Presence<br>(Median, IQR)<br>(Range) | HDQ Severity Score<br>(Median, IQR)<br>(Range) | HDQ Episodic<br>Presence Score<br>(Median; IQR)<br>(Range)* |
| --- | --- | --- | --- |
| Physical symptoms and<br>Impairments (20 items) | 25 (15, 45)<br>Range: 0-90 | 9 (4, 18)<br>Range: 0-58 | **5 (0, 20)**<br>**Range 0-80** |
| Cognitive symptoms and<br>impairments<br>(3 items) | 33 (0, 67)<br>Range: 0-100 | 8 (0,25)<br>Range: 0-100 | 0 (0, 0)<br>Range: 0-100 |
| Mental-emotional health<br>symptoms and impairments<br>(11 items) | 54 (27, 82)<br>Range: 0-100 | 18 (7, 34)<br>Range: 0-89 | 0 (0, 27)<br>Range: 0-100 |
| Uncertainty<br>(14 items) | **64 (43, 86)**<br>**Range: 0-100** | **27 (14, 41)**<br>**Range: 0-98** | 0 (0,7)<br>Range: 0-86 |
| Difficulties with Day-to-Day<br>Activities<br>(9 items) | 0 (0,22)<br>Range: 0-100 | 0 (0, 7)<br>Range: 0-61 | 0 (0,0)<br>Range: 0-89 |
| Challenges to Social Inclusion<br>(12 items) | 33 (17, 58)<br>Range: 0-100 | 12 (4, 27)<br>Range: 0-81 | 0 (0,0)<br>Range: 0-83 |
| Total HDQ Score | 38 (22, 57)<br>Range: 0-93 | 14 (8, 23)<br>Range: 0-70 | 2 (0, 16)<br>Range: 0-81 |

Higher scores indicate greater presence, severity and episodic nature of disability.
**Bold** indicates the highest score across all domains;
\*For the episodic scores, due to the higher rate of missingness we conducted a post hoc comparison and found no difference in episodic scores post median imputation.

### Criterion Measures

Similar to the HDQ, criterion measure summary scores were skewed to the healthier range of the scales (Shapiro Wilk Test for all criterion items and summary scores p<0.0001; data not shown). Median PHQ-9 scores were 4 out of possible range 0-27 (IQR: 2, 8) indicating ‘minimal depression severity. Median GAD scores were 10 out of possible range 0-21 (IQR: 8, 14) indicating low to moderate anxiety. Median scores of the international HIV dementia scale was 12 out of possible range 0-12 (IQR: 7,12), and 8 out of possible range 0-52 (IQR: 4, 15) on the EMQ, indicating high cognitive health. Median scores on the Wellness Thermometer was 7 out of possible range 0-10 (IQR: 5, 8) indicating participants reported to tend to feel well. Median scores of the Fatigue Scale were 29 out of possible range 9-63 (IQR: 21, 38) indicating participants may be approaching fatigue. Median scores on the ESS was 6 out of possible range 0-64 (IQR: 3, 9) indicating no evidence of abnormal daytime sleepiness in this sample.

### Reliability - Internal Consistency

#### HDQ Severity Scores

All individual items correlated with the HDQ Total Severity Score >0.2 except for Item #8 – ‘I have trouble swallowing food’ (r=0.14; p=0.03), and Item #15 – ‘I am unintentionally losing weight’ (r=0.19; p=0.03), and each item correlated with its corresponding domain score >0.20.

#### HDQ Episodic Scores

All individual items correlated with the HDQ Total Episodic Score >0.20 except for Item #8 (r=0.11; p=0.08) and Item #15 (r=0.15; p=0.02), and each item correlated with its corresponding domain score >0.20. Cronbach’s alpha for the entire HDQ was 0.96 (95% Confidence Interval (CI): 0.96-0.97) and ranged from 0.85 (95%CI: 0.80-0.90) in the cognitive domain to 0.93 (95% CI: 0.91-0.94) in the mental-emotional domain. The KR-20 statistic for the entire episodic scale of the HDQ was 0.95 (95% CI: 0.94-0.96) and ranged from 0.74 (95% CI: 0.66-0.83) in the cognitive domain to 0.91 (95% CI: 0.89-0.94) in the uncertainty domain (Table 3).

**Table 3.** Internal Consistency Reliability for HDQ Items (n=243)

| HDQ Items | HDQ Severity Scale |  | HDQ Episodic Scale |  |
| --- | --- | --- | --- | --- |
|  | Cronbach Alpha (Raw values) | 95% confidence interval | Kuder-Richardson Statistic (Raw values) | 95% confidence interval |
| HDQ Items (all) | 0.96 | 0.96, 0.97 | 0.95 | 0.94, 0.96 |
| Physical Symptoms and Impairments | 0.87 | 0.85, 0.90 | 0.84 | 0.80, 0.88 |
| Cognitive Symptoms and Impairments | 0.85 | 0.80, 0.90 | 0.74 | 0.66, 0.83 |
| Mental and Emotional Health Symptoms and Impairments | 0.93 | 0.91, 0.94 | 0.90 | 0.87, 0.92 |
| Uncertainty | 0.90 | 0.88, 0.92 | 0.91 | 0.89, 0.94 |
| Difficulty with Day-to-Day Activities | 0.90 | 0.86, 0.93 | 0.82 | 0.73, 0.91 |
| Challenges to Social Inclusion | 0.87 | 0.84, 0.90 | 0.84 | 0.79, 0.89 |

95% Confidence Interval: asymptotically distribution free (ADF) for non-normal data.
Median imputation of episodic scores; >0.8 defined as acceptable for individual patients

### Precision of Measurement

The standardised error of measurement (SEM) for HDQ items ranged from 0.05 (Item #8 – I have trouble swallowing food) to 0.28 (Item #64 – I find it hard to talk to others about my illness). Level of precision for the HDQ domain scores ranged from most precise in the difficulties with day-to-day activities domain (SEM: 3.71; SDC: 7.29) to the least precise in the cognitive symptoms domain (SEM: 7.68; SDC: 15.05) (Table 4).

**Table 4.**
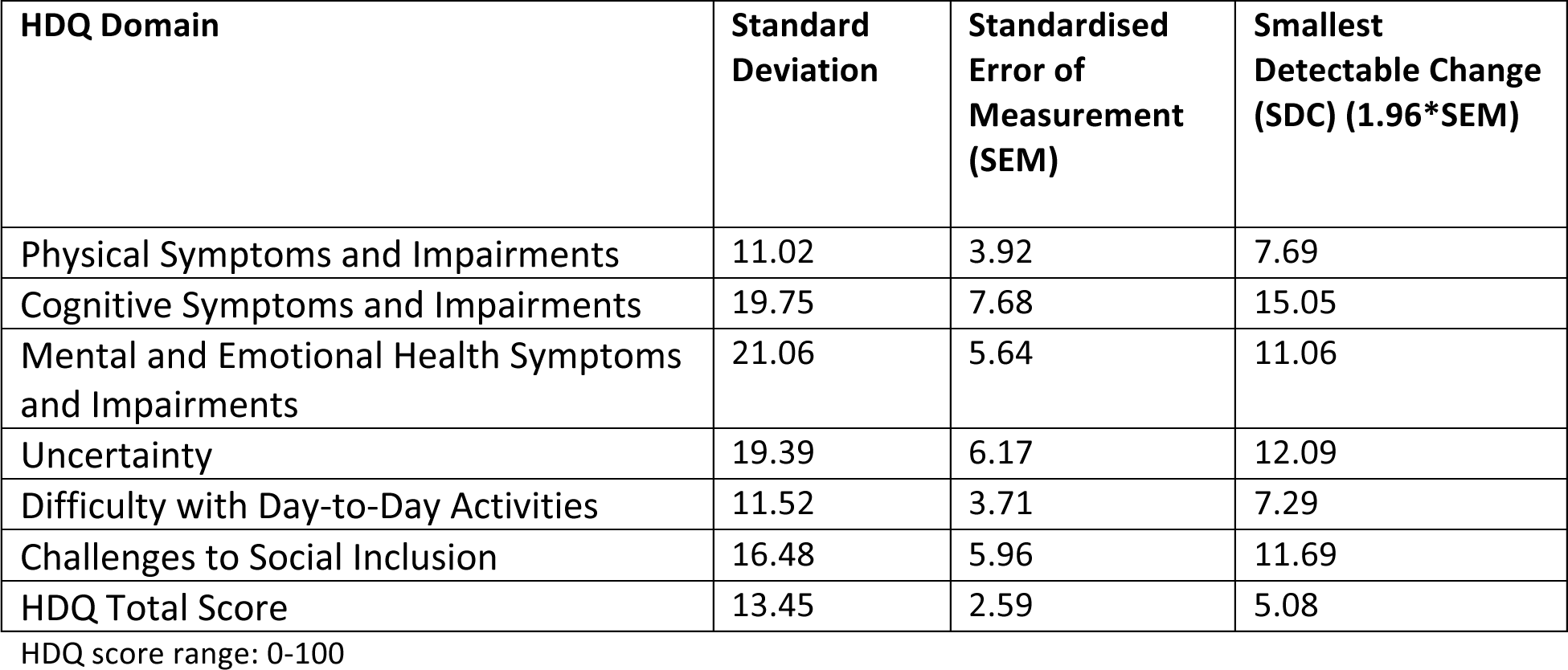
Level of Precision of HDQ Scores for Participants (n=243)

| HDQ Domain | Standard Deviation | Standardised Error of Measurement (SEM) | Smallest Detectable Change (SDC) (1.96*SEM) |
| --- | --- | --- | --- |
| Physical Symptoms and Impairments | 11.02 | 3.92 | 7.69 |
| Cognitive Symptoms and Impairments | 19.75 | 7.68 | 15.05 |
| Mental and Emotional Health Symptoms and Impairments | 21.06 | 5.64 | 11.06 |
| Uncertainty | 19.39 | 6.17 | 12.09 |
| Difficulty with Day-to-Day Activities | 11.52 | 3.71 | 7.29 |
| Challenges to Social Inclusion | 16.48 | 5.96 | 11.69 |
| HDQ Total Score | 13.45 | 2.59 | 5.08 |

HDQ score range: 0-100

### Construct Validity

Of the 36 construct validity hypotheses (8 primary, 28 exploratory), seven (88%) primary, 23 (28%) exploratory, and 30 (83%) of the total hypotheses were confirmed (Table 5).

**Table 5.** Construct Validity Analysis

| Construct Validity Analysis – *a priori* hypotheses | Spearman Correlation Coefficient (95% Confidence Interval) |
| --- | --- |
| **Convergent Construct Validity (22 hypotheses)** theorizing relationships between data collected in the HIV Disability Questionnaire (HDQ) and criterion measures |  |
| **PHQ-9** |  |
| 1) *Scores on PHQ-9 will be strongly correlated ($\geq 0.7$) with the mental and emotional symptoms domains of the HDQ. | **0.83 (0.63, 0.76)<sup>^</sup>** |
| 2) Scores on PHQ-9 will be moderately correlated ($\geq 0.5$) with the uncertainty domain of the HDQ. | **0.52 (0.37, 0.57)<sup>^</sup>** |
| 3) Scores on PHQ-9 will be moderately correlated ($\geq 0.5$) with the cognitive symptoms and impairments of the HDQ. | **0.66 (0.49, 0.66)<sup>^</sup>** |
| 4) Scores on PHQ-9 will be moderately correlated ($> 0.5$) with the challenges to social inclusion domain of the HDQ. | **0.62 (0.45, 0.63)<sup>^</sup>** |
| **General Anxiety Disorder (GAD) – 4 *a priori* hypotheses** |  |
| 5) *Scores on the General Anxiety Disorder (GAD) questionnaire will be strongly correlated ($\geq 0.7$) with the mental and emotional symptom domains of the HDQ. | **0.81 (0.59, 0.73)<sup>^</sup>** |
| 6) Scores on the General Anxiety Disorder (GAD) questionnaire will be moderately correlated ($> 0.5$) with the uncertainty domain of the HDQ. | **0.52 (0.37, 0.57)<sup>^</sup>** |
| 7) Scores on the General Anxiety Disorder (GAD) questionnaire will be | **0.56 (0.41, 0.60)<sup>^</sup>** |
| moderately correlated ($\geq 0.5$) with the cognitive symptoms and impairments domain of the HDQ. | |
| 8) Scores on the General Anxiety Disorder (GAD) questionnaire will be moderately correlated ($\geq 0.5$) with the challenges to social inclusion domain of the HDQ. | **0.61 (0.45, 0.63)^** |
| **Fatigue Scale** |  |
| 9) *Scores on the Fatigue Scale will be moderately correlated ($\geq 0.5$) with the physical symptoms domain of the HDQ. | **0.61 (0.45, 0.63)^** |
| 10) Scores on the Fatigue Scale will be moderately correlated ($\geq 0.5$) with the difficulties with day-to-day activity domain of the HDQ. | **0.57 (0.41, 0.60)^** |
| 11) Scores on the Fatigue Scale will be moderately correlated ($\geq 0.5$) with the challenges to social inclusion domain of the HDQ. | 0.39 (0.26, 0.48) |
| **Wellness Thermometer** |  |
| 12) *Scores on the Wellness Thermometer will be negatively moderately correlated ($\geq 0.5$) with the HDQ Total Score. | **-0.67 (-0.66, -0.50)^** |
| 13) Scores on the Wellness Thermometer will be negatively moderately correlated ($\geq 0.5$) with the PHYSICAL domain score on the HDQ. | **-0.64 (-0.65, -0.47)^** |
| 14) Scores on the Wellness Thermometer will be negatively moderately correlated ($\geq 0.5$) with the COGNITIVE domain score on the HDQ. | -0.41 (-0.49, -0.27) |
| 15) Scores on the Wellness Thermometer will be negatively moderately correlated ($\geq 0.5$) with the MENTAL-EMOTIONAL domain score on the HDQ. | **-0.69 (-0.67, -0.51)^** |
| 16) Scores on the Wellness Thermometer will be negatively moderately correlated ($\geq 0.5$) with the UNCERTAINTY domain score on the HDQ. | -0.43 (-0.51, -0.29) |
| 17) Scores on the Wellness Thermometer will be negatively moderately correlated ($\geq 0.5$) with the DIFFICULTIES WITH DAY-TO-DAY ACTIVITIES domain score on the HDQ. | **-0.50 (-0.56, -0.36)^** |
| 18) Scores on the Wellness Thermometer will be negatively moderately correlated ($\geq 0.5$) with the CHALLENGES TO SOCIAL INCLUSION domain score on the HDQ. | -0.49 (-0.55, -0.34) |
| **International Dementia Scale** |  |
| 19) *Scores on the International Dementia Scale (Total Score) will be strongly correlated ($\geq 0.7$) with the cognitive symptoms domain of the HDQ. | -0.09 (-0.21, 0.04) |
| **Everyday Memory Questionnaire** |  |
| 20) *Scores on the Everyday Memory Questionnaire (EMQ) will be strongly correlated ($\geq 0.7$) to the cognitive domain of the HDQ. | **0.73 (0.54, 0.67)^** |
| 21) Scores on the Everyday Memory Questionnaire will be moderately correlated ($> 0.5$) with the difficulties with day-to-day activity domain of the HDQ. | **0.54 (0.39, 0.58)^** |
| 22) Scores on the Everyday Memory Questionnaire will be moderately correlated ($> 0.5$) with the challenges to social inclusion domain of the HDQ. | 0.42 (0.29, 0.50) |
| **Divergent Construct Validity (7 hypotheses)** theorizing relationships between data collected in the HDQ and criterion measures |  |
| 23) *Scores on the Epworth Sleepiness Scale (ESS) will be weakly correlated ($\geq 0.30$) with the HDQ Total Score | **0.44 (0.30, 0.51)<sup>^</sup>** |
| 24) Scores on the Epworth Sleepiness Scale (ESS) will be weakly correlated ($\geq 0.30$) with PHYSICAL domain score on the HDQ. | **0.37 (0.23, 0.46)<sup>^</sup>** |
| 25) Scores on the Epworth Sleepiness Scale (ESS) will be weakly correlated ($\geq 0.30$) with COGNITIVE domain score on the HDQ. | **0.40 (0.26, 0.48)<sup>^</sup>** |
| 26) Scores on the Epworth Sleepiness Scale (ESS) will be weakly correlated ($\geq 0.30$) with MENTAL-EMOTIONAL domain score on the HDQ. | **0.34 (0.21, 0.44)<sup>^</sup>** |
| 27) Scores on the Epworth Sleepiness Scale (ESS) will be weakly correlated ($\geq 0.30$) with UNCERTAINTY domain score on the HDQ. | **0.32 (0.18, 0.42)<sup>^</sup>** |
| 28) Scores on the Epworth Sleepiness Scale (ESS) will be weakly correlated ($\geq 0.30$) with DIFFICULTIES WITH DAY-TO-DAY ACTIVITIES domain score on the HDQ. | **0.43 (0.29, 0.50)<sup>^</sup>** |
| 29) Scores on the Epworth Sleepiness Scale (ESS) will be weakly correlated ($\geq 0.30$) with CHALLENGES TO SOCIAL INCLUSION domain score on the HDQ. | **0.37 (0.24, 0.46)<sup>^</sup>** |
| **Known Groups Construct Validity (7 hypotheses)** theorizing relationships between data collected in the HDQ and Self-Perceived State of Health |  |
| *30-36) Participants who completed the HDQ on a 'good day' will have significantly lower scores on all HDQ domain scores and HDQ total scores; [7 hypotheses]<br>*\*HDQ Total was primary hypothesis* | **All 7 hypotheses confirmed# (p<0.001)<sup>^</sup>** |
| **Number of HDQ Construct Validity Hypotheses Confirmed** |  |
| **Primary Hypotheses** | 7/8 (88%) |
| **Exploratory Hypotheses** | 23/28 (82%) |
| **Total Hypotheses** | 30/36 (83%) |

\*Primary hypotheses; **Bold<sup>^</sup>** indicates significance (p<0.001); #Wilcoxon Test

## Discussion

The HDQ demonstrated internal consistency reliability and construct validity among a community dwelling sample of males living with HIV in an urban UK setting. Internal consistency reliability was achieved with Cronbach’s alpha and KR-20 statistics (scores >0.8) for all domain and total scores for episodic and severity scores, except for the cognitive domain for the episodic scale. This suggests that collectively items in the HDQ are homogenous within the six HDQ domains to collectively measure the broader construct of disability at one time point (33). Precision of measurement varied with subscales scores demonstrating highest levels of precision in the difficulties with day-to-day activities domain (SDC: 7.68), to lowest levels of precision in the cognitive symptoms domain (SDC: 15.05), suggesting among PLHIV, the HDQ possesses levels of measurement error and day-to-day variability. Construct validity was achieved as demonstrated by 88% primary (n = 7/8) and 83% total (n = 30/36) hypothesised relationships confirmed between the HDQ and criterion measures, which surpassed our 75% construct validity threshold (31). Our results build on previous evidence establishing internal consistency reliability and construct validity of the HDQ in Canada (32), Ireland (33), and the United States (34), as well as test-retest reliability in Canada (33).

Our study provides the first known assessment of HDQ psychometric properties including internal consistency reliability, construct validity, and level of precision of HDQ domain scores in the UK. Internal consistency reliability findings in this UK sample were similar to those among PLHIV for HDQ severity and episodic scores in Canada (α range: 0.87 - 0.97; KR-20 range: 0.81 - 0.98) (32), Ireland (α range: 0.84 - 0.96; KR-20 range: 0.85 - 0.96) (33), and the United States (α range: 0.89 - 0.93; KR-20 range: 0.87 - 0.96) (34), demonstrating the HDQ is reliable in measuring disability across high-income settings for PLHIV. Across all settings, Cronbach’s α or KR-20 were >0.8 for all domains and total scores for both episodic and severity disability scores, except for the UK cognitive symptoms and impairments domain for the episodic summary score (KR-20; 0.74). The cognitive domain possesses the fewest number of items (n=3), which might account for the lower alpha and KR-20 coefficients in this domain. Nevertheless, internal consistency reliability coefficients in this study exceeded the Special Advisory Committee of the Medical Outcomes Trust recommendations, that considers a Cronbach’s alpha of >0.70 to be acceptable (50).

Precision of the HDQ scores varied across HDQ domains ranging from a SDC of 7.68 (difficulties with day-to-day activities) to 15.50 (cognitive domain). The smaller the SDC, the more precise the domain. These values suggest the minimum difference in HDQ domain scores that would need to occur in order to be confident that an individual had a true change in disability beyond day-to-day variability or measurement error. Our study is the first to report on levels of precision of the HDQ. SEM dually reflects precision of an instrument, as well as the measure’s variation within a patient sample (51). Nevertheless, results are cross-sectional distribution based scores, and there is no universal consensus on how many SEMs an individual must change in order for a change in scores to be considered significant, nor clinically important (51). Future research should assess the interpretability of HDQ scores to determine the meaning of HDQ scores (cross-sectionally) as well as the minimally clinically importance difference (MCID) (longitudinally) that represent the important ‘amount’ and ‘importance’ of change in disability over time.

The HDQ possesses construct validity in this UK sample, for its ability to measure disability as demonstrated by confirmation of total hypothesised relationships between HDQ and criterion measures (83%), which was above our *a priori* defined threshold of 75% (31). Construct validity was similarly demonstrated in Canada (80%) (33) and the United States (87%) (34), and also was demonstrated in Canada using confirmatory factor analysis (32). However it is not possible to compare the UK construct validity results to these previous studies, because the UK analysis used different criterion measures.

While the HDQ overall demonstrated internal consistency reliability and construct validity for use among males with HIV in the UK and PLHIV in other high income countries, reasons may exist for variations in HDQ scores and properties across different cultural contexts. Diversity in sample populations, recruitment procedures, and mechanisms in which the HDQ and reference measures were administered, may account for differences in HDQ scores and measurement property coefficients. For instance, UK participants were all male, mostly economically active and university educated, and living with well controlled HIV recruited from an HIV clinic setting compared with HDQ assessment with PLHIV in Ireland (33), where fewer participants were working for pay, and had been living longer with their HIV diagnosis. Moreover participants in Canada (32) were older, living with more comorbidities, and fewer working for pay, when recruited from community-based organisations. Furthermore, UK participants completed measurements either during their clinic visit or independently at home following their routine outpatient HIV care appointments, while Irish participants completed measures intermittently while seeing various health providers in a busy HIV outpatient setting, and Canadian participants completed measures consecutively in one single sitting in a quiet location at an HIV service organisation (33). This may have introduced inconsistencies in the way participants responded to items across the questionnaires, creating variations in correlations between measures (33). Similarly the different criterion measures used in the UK analysis may have resulted in different estimations of the extent to which we hypothesised items in the HDQ would correlate with items included in these criterion measures. Notably, our UK analysis did not include universal measures of disability, therefore to compare to other conditions a generic disability measurement tool might be recommended (e.g. World Health Organization Disability Assessment Schedule 2.0) (52). Hence, measurement properties should be interpreted cautiously and specific to the context and sample population.

Our results indicate that the HDQ domain of uncertainty or worrying about the future, was the most present and severe domain of disability in this UK sample of PLHIV. Uncertainty is a unique domain of disability within the *Episodic Disability Framework* (53). It is also a core dimension of disability experienced by adults ageing with HIV (54). Older PLHIV may worry about HIV specific age-related uncertainties (55) and the trajectory of episodic disability (56). The role of uncertainty has also been incorporated into rehabilitation recommendations for adults ageing with HIV (57), whereby interventions can promote stability, mitigate increasing disability, and increase time between episodes (56). Our results indicate that uncertainty can be experienced across the life-course among a younger sample of PLHIV in the UK, building on existing literature that uncertainty is the most present and severe domain of disability experienced by PLHIV in Canada (32), Ireland (33), and the United States (34). In this UK sample, the most episodic domain of disability was in the physical domain, which was similarly observed in Ireland (33) and the United States (34). This is likely attributed to health challenges in this domain more likely to fluctuate on a daily basis (e.g. aches and pains, fatigue) opposed to items in the social inclusion domain (e.g. employment, relationships), which may fluctuate over a longer duration of time. Further exploration is warranted into the experiences of uncertainty and episodic health challenges across the life-course among PLHIV in the UK, and the impact of rehabilitation, such as group-based interventions (58) to address disability including uncertainty.

Our study has limitations. Firstly participants were all male, living in an urban setting, therefore this sample is not representative of the UK population of PLHIV who are 69% male and 36% living in London (4). The HDQ was developed primarily with men living with HIV in a large metropolitan city, which may explain the high construct validity in this study, as this study sample might resemble the sample from which the HDQ was originally derived, validated, and refined in Ontario, Canada (33). Nevertheless, evaluation of the psychometric properties of the HDQ in other low to middle income contexts is warranted. Secondly, given this study was part of a larger cohort study (IRAS 165402), the criterion measures to assess construct validity were not consistent with previous HDQ psychometric evaluations (32-34). Therefore caution should be applied when comparing the validity and reliability of the HDQ. Next, because our goal was to assess the measurement properties of the HDQ in the UK, rather than to measure disability experienced by PLHIV in the UK, HDQ scores should be interpreted cautiously. Lastly, our analysis assessed internal consistency reliability and construct validity of the HDQ in the UK. Further analysis of the reproducibility, responsiveness and interpretability of the HDQ among PLHIV in the UK is needed.

Identification of the HDQ as a valid and reliable self-reported disability assessment tool has important implications for clinical practice, research and policy. Clinicians, HIV community organisations, and researchers, may use the HDQ to assess disability experienced by PLHIV in the UK. To our knowledge, there is no known evidence exploring disability experienced by PLHIV in the UK. Data on disability experienced by PLHIV, capturing multiple domains of functional limitations, is therefore required in the UK. Measuring disability can provide information on the nature and extent of disability, and the health care needs of PLHIV in the UK. This knowledge can help to inform ways in which HIV services can adopt approaches to better respond to the changing needs of PLHIV (23), while ensuring function is incorporated into the provision of person-centered care (59). Results provide a foundation for future research to utilise the HDQ to examine the extent and nature of disability among PLHIV in the UK and international cross-cultural comparisons of disability for PLHIV.

## Conclusions

The HDQ possesses internal consistency reliability and construct validity with varied levels of precision across domain scores, when administered to adults living with HIV in the UK. Results are specific to a mainly community dwelling sample of males, who are mostly economically active and university educated, living with well-controlled HIV. Future research should examine HDQ properties in low-to middle income countries, responsiveness to change, and interpretability of HDQ scores. Future research should consider cross-cultural, international comparisons of disability, and the ability of the HDQ to detect clinically important changes in disability for examining effectiveness of interventions.

## Acknowledgments

We gratefully acknowledge the support of St Stephen’s Clinical Research in funding and managing the Dean Street Cohort Study. We gratefully acknowledge the Dean Street Cohort Study participants, involved in this study. We also gratefully acknowledge the British Academy Fellowship of Humanities and Social Sciences (Harding, O’Brien), which supported this study. Kelly O’Brien is supported by a Canada Research Chair in Episodic Disability and Rehabilitation.

